# Resiniferatoxin hampers the nocifensive response of *Caenorhabditis elegans* to noxious heat, and pathway analysis revealed that the Wnt signaling pathway is involved

**DOI:** 10.1101/2021.07.26.453516

**Authors:** Jennifer Ben Salem, Bruno Nkambeu, Dina N Arvanitis, Francis Beaudry

## Abstract

Resiniferatoxin (RTX) is a metabolite extracted from *Euphorbia resinifera*. RTX is a potent capsaicin analog with specific biological activities resulting from its agonist activity with the transient receptor potential channel vanilloid subfamily member 1 (TRPV1). RTX has been examined as a pain reliever, and more recently, investigated for its ability to desensitize cardiac sensory fibers expressing TRPV1 to improve chronic heart failure (CHF) outcomes using validated animal models. *Caenorhabditis elegans* (*C. elegans*) expresses orthologs of vanilloid receptors activated by capsaicin, producing antinociceptive effects. Thus, we used *C. elegans* to characterize the antinociceptive properties and performed proteomic profiling to uncover specific signaling networks. After exposure to RTX, wild-type (N2) and mutant *C. elegans* were placed on petri dishes divided into quadrants for heat stimulation. The thermal avoidance index was used to phenotype each tested *C. elegans* experimental group. The data revealed for the first time that RTX can hamper the nocifensive response of *C. elegans* to noxious heat (32°C – 35°C). The effect was reversed 6 h after RTX exposure. Additionally, we identified the RTX target, the *C. elegans* transient receptor potential channel OCR-3. The proteomics and pathway enrichment analysis results suggest that Wnt signaling is triggered by the agonistic effects of RTX on *C. elegans* vanilloid receptors.

## Introduction

Resiniferatoxin (RTX) is a capsaicin analog originating from a Euphorbia species (*Euphorbia resinifera*) that targets the transient receptor potential channel vanilloid subfamily member 1 (TRPV1) and is several-fold more pungent than capsaicin [1]. One major pain relief strategy that has been tested in recent years includes pharmacological manipulation of TRPV1 [19, 20, 22–24]. However, TRPV1 activation by high doses of certain vanilloids can lead to necrosis and apoptosis [25]. RTX has been tested for chronic pain management [2] and cancer pain [3]. Recently, RTX was used to perform cardiac sympathetic afferent denervation, and the results showed that it reduces cardiac remodeling and improves cardiovascular function in a validated rat model of heart failure [4]. Chemical denervation results from the agonistic effects of RTX on TRPV1, producing an intense pungent effect and leading to cell death. Other cardiovascular benefits were observed, including a reduction in ventricular arrhythmias [5] and the lowering diastolic blood pressure [6]. Recently, we demonstrated that cardiac sensory afferent denervation by RTX during myocardial infarction could modulate long-term susceptibility to anxiodepressive behavior using a validated mouse model of chronic heart failure [7]. Treatment with RTX triggered the desensitization of cardiac afferents and hampered the anxiodepressive-like state in mice. Depression and anxiety are widely observed in patients suffering chronic heart failure [41]. Additionally, we found unique protein profiles and regulatory pathways in the frontal cortices that are associated with the response to stresses originating from the heart. RTX treatment produces unique brain signaling patterns that require further characterization. To extend our knowledge, we intend to use a model organism for which a large proportion of its genes have functional counterparts in mammals, which therefore makes these models exceptionally useful to study biological processes and diseases in mammals, including humans [26].

*Caenorhabditis elegans* (*C. elegans*) is a very interesting model organism, particularly in the context of functional genomics relevant to mammalian and human biology and diseases [8, 9, 26]. *C. elegans* genome sequencing was completed in 1998 and is publicly available and well annotated. This is a significant benefit of proteomics, as it enables the interpretation of functionally grouped gene ontology (GO) and pathway annotation networks [10]. There are roughly ~21,000 predicted protein-coding genes in *C. elegans*. Notably, many of these genes are shared with mammals, including humans [11, 12]. *C. elegans* adults consist of 959 cells, of which 302 are neurons, making this model a valuable model organism to study the nervous system [15]. Thus, *C. elegans* is particularly useful to study nociception, as the animal displays well-defined and reproducible nocifensive behavior involving a reversal and change in direction away from noxious stimuli. In mammals, important molecular transducers of noxious stimuli include ligand-gated (e.g., TRPV1, TRPA1) and voltage-gated (Nav1.7, Nav1.8, Nav1.9) ion channels [13]. Following *C. elegans* genome sequencing, several genes encoding TRP ion channels with important sequence homologies to mammalian TRP channels, including TRPVs, were identified [14]. More specifically, five TRPV orthologs (e.g., OSM-9, OCR-1, OCR-2, OCR-3 and OCR-4) were discovered and studied [16]. Furthermore, several studies have shown that *C. elegans* TRPV orthologs are associated with behavioral and physiological processes, including sensory transduction [17–20]. Additionally, it was determined that *C. elegans* TRPV channels share similar activation and regulatory mechanisms with higher species, including mammals. Interestingly, our recent paper revealed that capsaicin can impede the nocifensive response of *C. elegans* to noxious heat (i.e., 32°C – 35°C) following sustained exposure, and this effect was reversed 6 h after capsaicin exposure [19]. Furthermore, data suggest that capsaicin targets the *C. elegans* transient receptor potential ion channel OCR-2. Additional experiments indicated that other capsaicin analogs have antinociceptive effects, including olvanil, gingerol, shogaol and curcumin, as well as for other vanilloids, including eugenol, vanillin and zingerone [20]. Thus, known TRPV ligands that target *C. elegans* vanilloid receptors and produce antinociceptive effects compatible with previous studies were performed using animal models of pain.

Our hypothesis is that RTX will interact with TRPV ortholog channels in *C. elegans* in a manner similar to that of capsaicin, and sustained exposure to RTX will hamper the nocifensive response of *C. elegans* to noxious heat. The objective of this study was to 1) characterize the RTX exposure–response relationships using *C. elegans* and heat avoidance behavior analysis [21] and 2) perform proteomics to identify the proteins and pathways responsible for the induced phenotype.

## Materials and methods

### Chemicals and reagents

All chemicals and reagents were obtained from Fisher Scientific (Fair Lawn, NJ, USA) or Millipore Sigma (St. Louis, MO, USA). Capsaicin was purchased from Toronto Research Chemicals (North York, ON, CAN). RTX was obtained from Tocris Bioscience (Rennes, France).

### *C. elegans* strains

The N2 (Bristol) isolate of *C. elegans* was used as a reference strain. The mutant strains used in this study included *ocr-1* (ak46), *ocr-2* (yz5), *ocr-3* (ok1559)*, ocr-4* (vs137) and *osm-9* (yz6). N2 (Bristol) and other strains were obtained from the Caenorhabditis Genetics Center (CGC), University of Minnesota (Minneapolis, MN, USA). Strains were maintained and manipulated under standard conditions as previously described [27, 28]. Nematodes were grown and kept on nematode growth medium (NGM) agar at 22°C in a Thermo Scientific Heratherm refrigerated incubator (Fair Lawn, NJ, USA). Analyses were executed at room temperature (~22°C) unless otherwise noted.

### *C. elegans* pharmacological manipulations

Capsaicin or RTX was dissolved in Type 1 Ultrapure Water at a concentration of 25 μM. The solutions were warmed for brief periods combined with vortexing and sonication for several minutes to completely dissolve the compounds. Further dilutions to concentrations of 5 μM, 1 μM and 0.1 μM in Type 1 Ultrapure Water were performed by serial dilution of the RTX stock solution. *C. elegans* was isolated and washed according to the protocol outlined by Margie *et al*. [28]. After 72 h of feeding and growing on 92 × 16 mm petri dishes with NGM, the nematodes were denied food and exposed to capsaicin or RTX solutions. An aliquot of 7 mL of a capsaicin or RTX solution was added to produce a 2-3 mm solution film (the solution was partly absorbed by NGM); consequently, the nematodes swam in solution. *C. elegans* was exposed to capsaicin or RTX for 60 min and isolated and washed thoroughly prior to the behavioral experiments. For the residual effect (i.e., 6 h latency) evaluation, after exposure to the RTX solutions, the nematodes were isolated, carefully washed and deposited on NGM free of RTX for 6 h prior to testing.

### Thermal avoidance assays

The method we proposed in this manuscript for the evaluation of thermal avoidance was modified from the four quadrant strategy previously described [29] and used in previous successfully published work [19–21]. Briefly, experiments were performed on 92 × 16 mm petri dishes divided into four quadrants. A middle circle delimited (i.e., 1 cm diameter) an area where *C. elegans* were not considered. Petri dishes were divided into quadrants: two stimulus areas (A and D) and two control areas (B and C). Sodium azide (0.5 M) was used in all quadrants to paralyze the nematodes. Noxious heat was created with an electronically heated metal tip (0.8 mm in diameter) producing a radial temperature gradient (e.g., 32-35°C on NGM agar 2 mm from the tip measured with an infrared thermometer). Nematodes were isolated and washed according to a protocol outlined by Margie *et al.* [28]. The nematodes tested were denied food during all experimentations. The nematodes (typically 100 to 1,000 young adult nematodes) were placed at the center of a marked petri dish, and after 30 min, the plates were placed at 4°C for at least 1 h, and the number of nematodes per quadrant were counted. Nematodes that did not cross the inner circle were not considered. The thermal avoidance index (TI) was calculated as shown in equation 1.

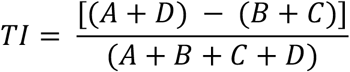

Details are shown in supplementary Figure S1. Both the TI and the animal avoidance percentage were used to phenotype each tested *C. elegans* experimental group. The stimulus temperature used was based on previous experiments [30].

### Proteomic analysis

Cultured nematodes (with or without RTX exposure) were collected in liquid media and centrifuged at 1,000 g for 10 min; the nematodes were then collected and thoroughly washed. Nematodes were resuspended in 8 M urea in 100 mM TRIS-HCl buffer (pH 8) containing cOmplete™ protease inhibitor cocktails (Roche Diagnostic Canada, Laval, QC, Canada) and aliquoted into reinforced 1.5 mL homogenizer tubes containing 500 μm glass bead homogenizer tubes with 25 mg of glass beads. The samples were homogenized using a Bead Mill Homogenizer (Fisher) with 3 bursts of 60 seconds at a speed of 5 m/s. The homogenates were centrifuged at 9,000 g for 10 min. The protein concentration for each homogenate was determined using a Bradford assay. Two hundred micrograms of protein was extracted using ice-cold acetone precipitation (1:5, v/v). The protein pellet was dissolved in 100 μL of 50 mM TRIS-HCl buffer (pH 8), and the solution was mixed with a Disruptor Genie at maximum speed (2,800 rpm) for 15 min and sonicated to improve the protein dissolution yield. The proteins were denatured by heating at 120°C for 10 min using a heated reaction block. The solution was allowed to cool 15 min. Proteins were reduced with 20 mM dithiothreitol (DTT), and the reaction was performed at 90°C for 15 min. Then, proteins were alkylated with 40 mM iodoacetamide (IAA protected from light at room temperature for 30 min. Then, 5 μg of proteomic-grade trypsin was added, and the reaction was performed at 37°C for 24 h. Protein digestion was quenched by adding 10 μL of a 1% trifluoroacetic acid (TFA) solution. Samples were centrifuged at 12,000 g for 10 min, and 100 μL of the supernatant was transferred into injection vials for analysis. The HPLC system was a Thermo Scientific Vanquish FLEX UHPLC system (San Jose, CA, USA). Chromatography was performed using gradient elution along with a Thermo Biobasic C18 microbore column (150 × 1 mm) with a particle size of 5 μm. The initial mobile phase conditions consisted of acetonitrile and water (both fortified with 0.1% formic acid) at a ratio of 5:95. From 0 to 3 min, the ratio was maintained at 5:95. From 3 to 123 min, a linear gradient was applied up to a ratio of 40:60, which was maintained for 3 min. The mobile phase composition ratio was the reverted to the initial conditions, and the column was allowed to re-equilibrate for 30 min. The flow rate was fixed at 50 μL/min, and 5 μL of each sample was injected. A Thermo Scientific Q Exactive Plus Orbitrap Mass Spectrometer (San Jose, CA, USA) was interfaced with the UHPLC system using a pneumatic-assisted heated electrospray ion source. Nitrogen was used as the sheath and auxiliary gases, which were set at 15 and 5 arbitrary units, respectively. The auxiliary gas was heated to 200°C. The heated ESI probe was set to 4000 V, and the ion transfer tube temperature was set to 300°C. Mass spectrometry (MS) detection was performed in positive ion mode operating in TOP-10 data dependent acquisition (DDA) mode. A DDA cycle entailed one MS^1^ survey scan (m/z 400-1500) acquired at 70,000 resolution (FWHM) and precursor ions meeting the user-defined criteria for charge state (i.e., z = 2, 3 or 4), monoisotopic precursor intensity (dynamic acquisition of MS^2^-based TOP-10 most intense ions with a minimum 1×10^4^ intensity threshold). Precursor ions were isolated using the quadrupole (1.5 Da isolation width), activated by HCD (28 NCE) and fragment ions were detected in the ORBITRAP at a resolution of 17,500 (FWHM). Data were processed using Thermo Proteome Discoverer (version 2.4) in conjunction with SEQUEST using default settings unless otherwise specified. The identification of peptides and proteins with SEQUEST was performed based on the reference proteome extracted from UniProt (*C. elegans* taxon identifier 6239) as FASTA sequences. Parameters were set as follows: MS^1^ tolerance of 10 ppm; MS^2^ mass tolerance of 0.02 Da for Orbitrap detection; enzyme specificity was set as trypsin with two missed cleavages allowed; carbamidomethylation of cysteine was set as a fixed modification; and oxidation of methionine was set as a variable modification. The minimum peptide length was set to six amino acids. Data sets were further analyzed with Percolator. Peptide-spectrum matches (PSMs) and protein identification were filtered at a 1% false discovery rate (FDR) threshold. For protein quantification and comparative analysis, we used the peak integration feature of Proteome Discoverer 2.4 software. For each identified protein, the average ion intensity of the unique peptides was used for protein abundance.

### Bioinformatics

Abundance Ratio (log2): (experimental group)/(N2 control), Abundance Ratio P-Value: (experimental group)/(N2 control) and Accession columns were extracted from the datasets generated by Proteome Discoverer. Volcano plots were generated using all identified and quantified proteins with both a log2 ratio and p-value. Proteins without a p-value or with a p-value > 0.01 were not used for further functional analysis. Additionally, only proteins with an absolute log_2_ ratio ≥ 1.0 were used for bioinformatics analysis. Gene Ontology analysis of genes (i.e., GO terms) was performed using Metascape [31], where two sets of proteins (e.g., Ensembl gene codes) were used: proteins with a log_2_ ratio ≥ 1.0 (i.e., upregulated) and proteins with a log_2_ ratio ≤ −1.0 (downregulated). The results from the upregulated and downregulated protein analyses were used to create bar plots in PRISM using the Metascape reports with the - log_10_ p-value of GO process matches given by Metascape. Further pathway analysis was performed using ClueGO [32] and Cytoscape [33] using the KEGG and REACTOME pathway databases. Significant pathways for proteins with an absolute log_2_ ratio ≥ 1.0 were identified, comparing the ratio of target genes identified in each pathway to the total number of genes within the pathway. The statistical test used to determine the enrichment score for KEGG or REACTOME pathways was based on a right-sided hypergeometric distribution with multiple testing correction (Benjamini & Hochberg (BH) method) [34].

### Statistical analysis

Behavioral data were analyzed using one-way ANOVA followed by Dunnett’s multiple comparison test (e.g., WT (N2) was the control group) or ANOVA followed by a Tukey-Kramer multiple comparison test. Data presented in Figure 4C were analyzed using ANOVA followed by Sidak’s multiple comparisons test for specific pairwise comparisons. Significance was set a priori to p < 0.05. Statistical analyses were performed using PRISM (version 9.1.2).

## Results and discussion

Recently, we demonstrated for the first time the antinociceptive effects of capsaicin in *C. elegans* following controlled and prolonged exposure [19]. Further analysis revealed that capsaicin targets OCR-2, one of the five vanilloid receptor orthologs (e.g., OSM-9, OCR-1, OCR-2, OCR-3 and OCR-4). Moreover, we have shown that other vanilloids, including eugenol, induce similar antinociceptive effects [20]. The thermal avoidance assay performed is described in supplementary Figure S1 and was specifically used to assess the antinociceptive effects of RTX. The first step was to verify whether there was bias in *C. elegans* behavior using the experimental setup. Thus, we performed an assessment of the mobility and bias of WT (N2) nematodes with and without exposure to RTX. As shown in Figure 1, no quadrant selection bias was observed for all *C. elegans* experimental groups tested with or without RTX exposure (e.g., 5, 1 and 0.1 μM). The data demonstrated that the nematodes did not select any specific quadrant and were uniformly distributed 30 min after the initial nematode deposition at the center of the petri dish. Sixty minutes of RTX exposure did not appear to affect nematode mobility. In a previous study, we tested mutant nematodes *ocr-1*, *ocr-2*, *ocr-3, ocr-4* and *osm-9* [19, 20], and no bias was observed.

**Figure 1.**
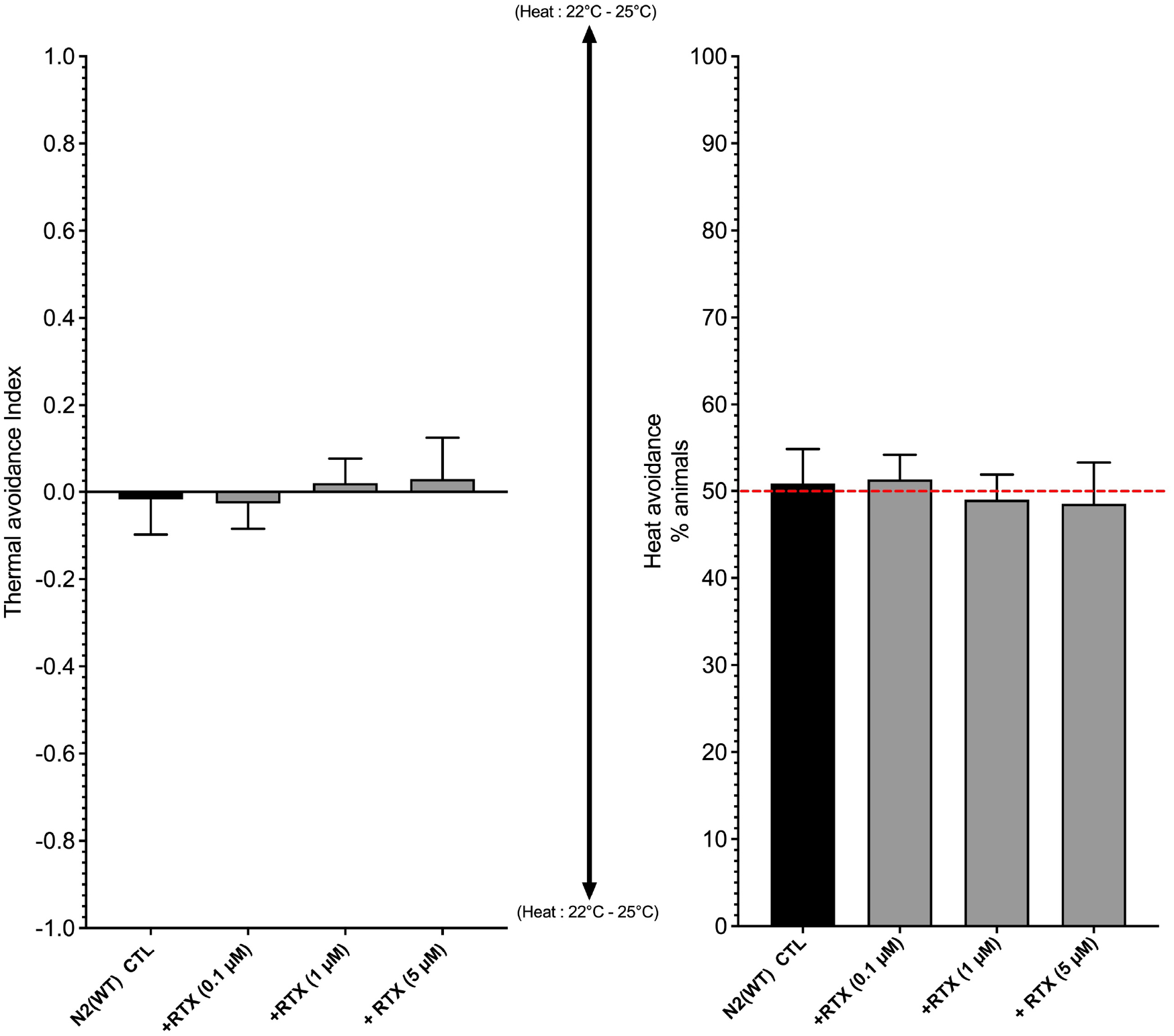
Comparison of the mobility and bias of WT (N2) nematodes in plates divided into quadrants conserved at a constant temperature (22°C) without the application of a stimulus (negative control). No quadrant selection bias was observed for any *C. elegans* genotype tested in the absence or presence of RTX at 5 μM, 1 μM and 0.1 μM.

Prolonged stimulation with TRPV1 agonists elicits receptor desensitization, leading to the alleviation of pain (or antinociceptive effects in *C. elegans*) [22]. Consequently, we exposed *C. elegans* to RTX in solution for the complete control of time and exposure levels. As presented in Figure 2, the data revealed antinociceptive effects following a 1 h of exposure, which were, to a certain extent concentration-dependent. The antinociceptive effects of RTX were noted even at a very low concentration (0.1 μM), and RTX was found to be significantly more potent than capsaicin [19]. These observations are consistent with previously published in vitro and in vivo testing data [35]. Following RTX exposure, the nematodes were carefully washed and transferred to NGM agar maintained at 22°C in an incubator for 6 h (i.e., residual effect/latency test). Then, the thermal avoidance response was retested. As shown in Figure 2, the results indicated that 6 h after exposure to RTX, the *C. elegans* thermal avoidance response returned to normal, confirming that no residual antinociceptive effects of RTX were observed after 6 h. These results were unexpected and are quite interesting since RTX is apoptotic [25, 38] and the local application of RTX has been used for chemical denervation [7, 39]. The results suggest that RTX does not induce a long-lasting effect. Additionally, the results indicate that RTX does not induce significant damage to the sensory system at a concentration ≤ 5 μM and a 60-min exposure period.

**Figure 2.**
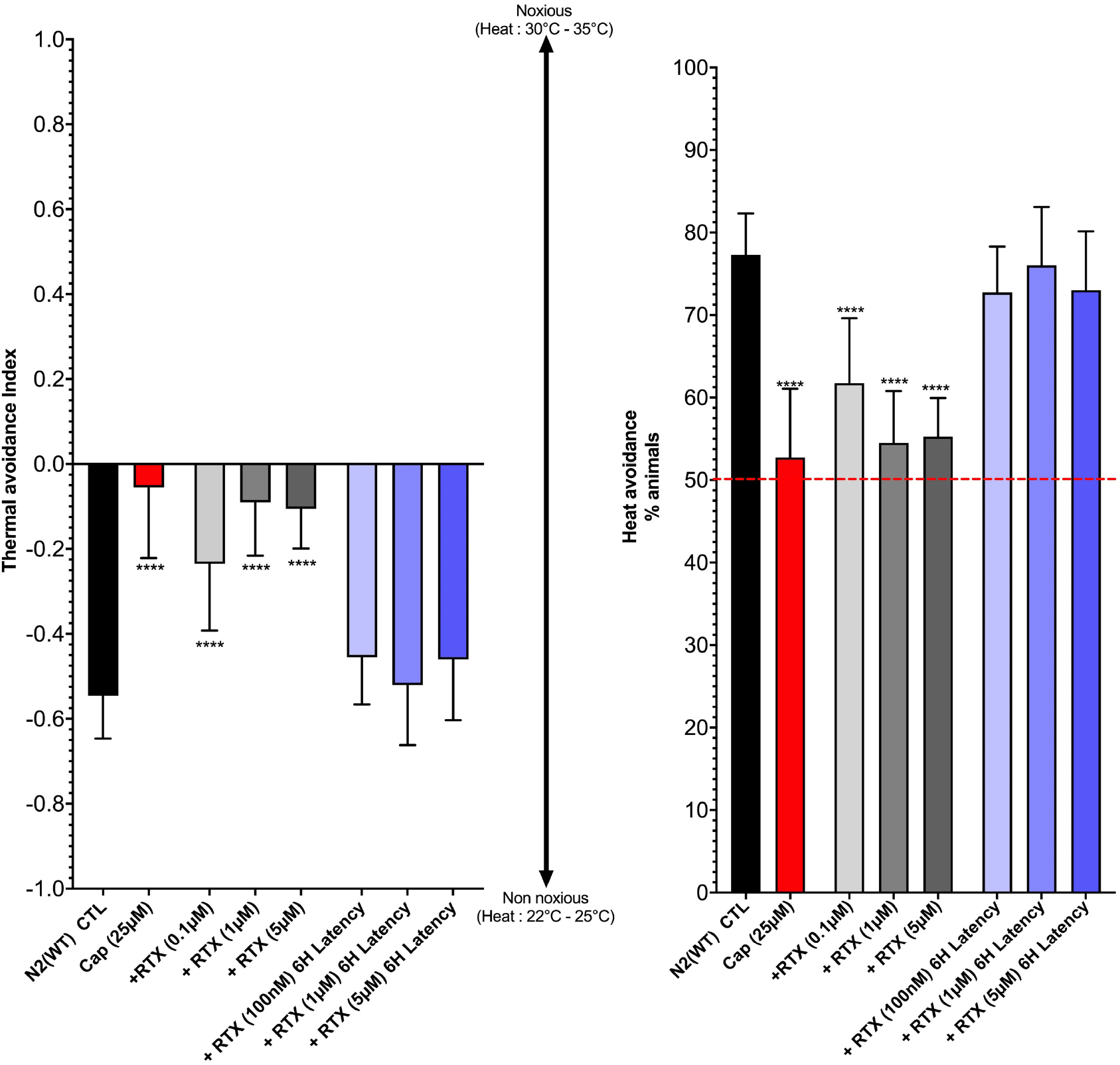
Assessment of the pharmacological effects of RTX on thermal avoidance in *C. elegans*. Nematodes were exposed to RTX or capsaicin (Cap) for 60 min prior to behavior experimentation. Display values (means ± SD) were calculated from at least 12 independent experiments for each experimental group. The observed effects of RTX are dose-dependent and noticeably hamper thermal avoidance in *C. elegans*. **** p < 0.0001 (ANOVA - Tukey-Kramer multiple comparison test). The results suggest an effect similar to that of capsaicin but at a notably lower concentration. No residual antinociceptive effects were observed 6 h after exposure (ANOVA - Dunnett’s multiple comparison versus CTL group).

The target of RTX needs to be identified to better understand the exposure–response relationship. Pharmacological effects can be measured when the tested molecule binds to a sufficient fraction of the target receptor, leading to a specific response (i.e., physiological or phenotypic changes). Accordingly, we conducted experiments using specific *C. elegans* mutants (*ocr-1*, *ocr-2*, *ocr-3*, *ocr-4* and *osm-9*) to potentially identify the RTX target receptors. *C. elegans* mutants were exposed to RTX at a concentration of 1 μM for 60 min prior to the behavioral experiments. As revealed in Figure 3, all mutants appeared less sensitive to noxious heat than the WT (N2) group. However, they were still sensitive, which suggests redundancy in receptor function. These results are compatible with our previous findings [19, 20]. Furthermore, Figure 3 shows that the RTX antinociceptive effects were quantifiable in the *ocr-1*, *ocr-2*, *ocr-4* and *osm-9* mutants. However, no significant RTX effects (p > 0.05) were observed with the *ocr-3* mutant, suggesting that RTX targets ORC-3, a mammalian vanilloid receptor-like channel. These results were unexpected since our previous results clearly showed that capsaicin targets the OCR-2 receptor, not OCR-3 [19]. Additionally, we showed that other vanilloids target OCR-2 [20]. As shown by another study, heat avoidance is mediated by OSM-9 and OCR-2 in *C. elegans* [17]. The functions of OCR-3 are unknown, but it is coexpressed with OSM-9 [40]. There is no clear evidence that OCR-3 is activated by nociceptive heat, but it is predicted to enable ion channel activity and is classified as capsaicin receptor-related. Little is known about this vanilloid receptor, and further study is required to elucidate OCR-3 function in *C. elegans*.

**Figure 3.**
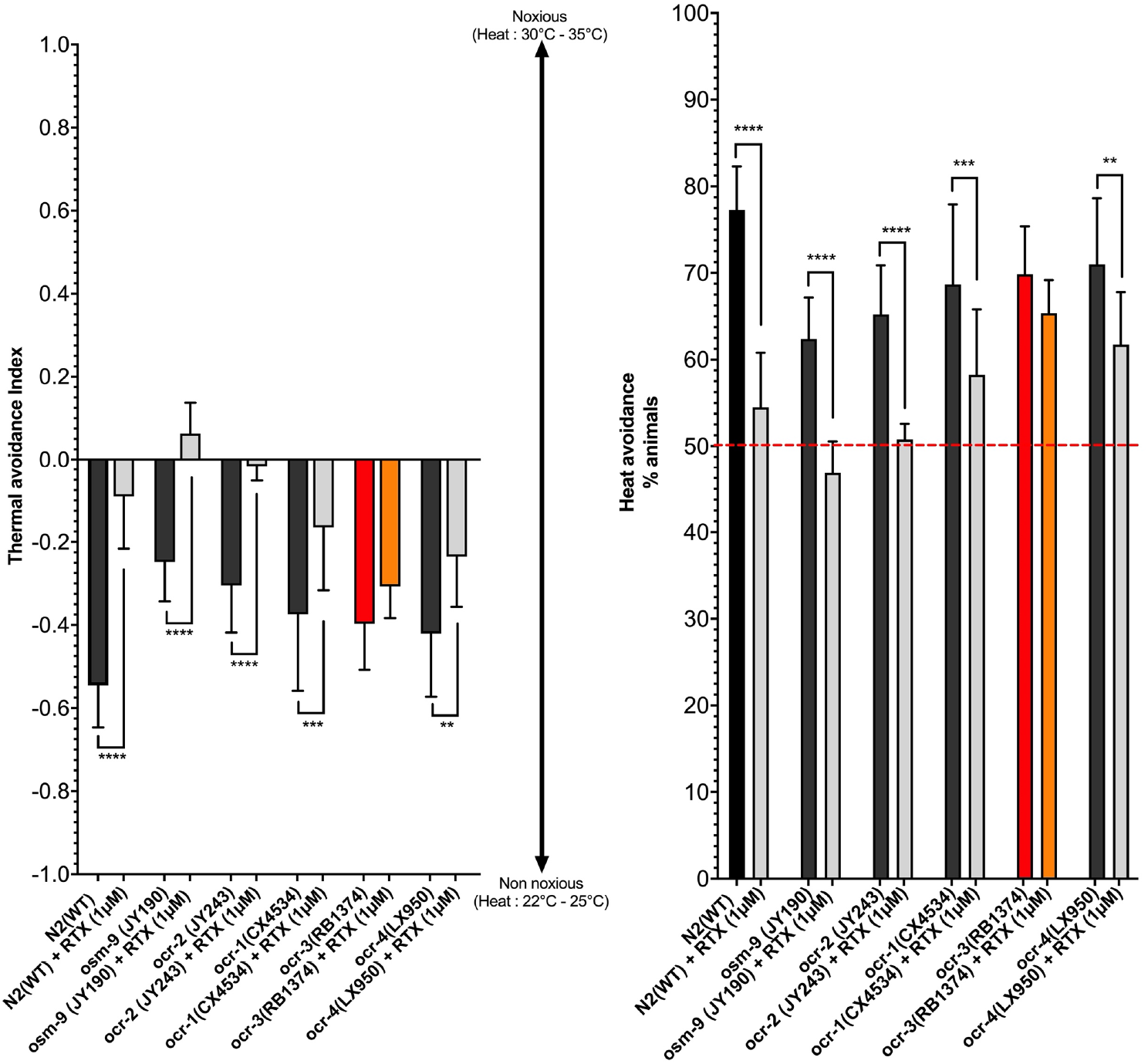
Identification of TRPV orthologs responsible for the RTX-induced antinociceptive effects. The data strongly suggest that capsaicin exerts antinociceptive effects through the ORC-3 *C. elegans* TRPV ortholog. **** p < 0.0001, *** p < 0.001, ** p < 0.01 (ANOVA - Sidak’s multiple comparisons test using specific comparison pairs).

Label-free proteomic investigations were performed on *C. elegans* exposed to 1 μM RTX for 60 min. Figure 4 shows volcano plots illustrating the differential abundances of proteins, with the x-axis representing the log_2_ ratio and the y-axis plotting −log_10_ (P-value). The colored boxes represent a 2-fold change and p-value ≤ 0.05. Several differentially expressed proteins (DEPs) were identified, including 157 upregulated and 305 downregulated proteins. Moreover, we extracted specific data related to vanilloid receptor proteins (i.e., OCR-1, OCR-2, OCR-3, OCR-4 and OSM-9) and the NPR-1 signaling pathway [17, 21]. Interestingly, OCR-1, OCR-2, OCR-3, OCR-4, OSM-9, FLP-18 and FLP-21 were significantly upregulated. Both FPL-18 and FPL-21 are important neuropeptides that interact with NPR-1 and are involved in heat sensory processes [17, 21]. Vanilloid receptors are mainly heat thermosensors and calcium-selective channels. Upregulation of vanilloid receptors can increase neuron depolarization, leading to receptor desensitization and conformational changes. These are major mechanisms that lead to the alleviation of pain or antinociceptive effects in *C. elegans*.

**Figure 4.**
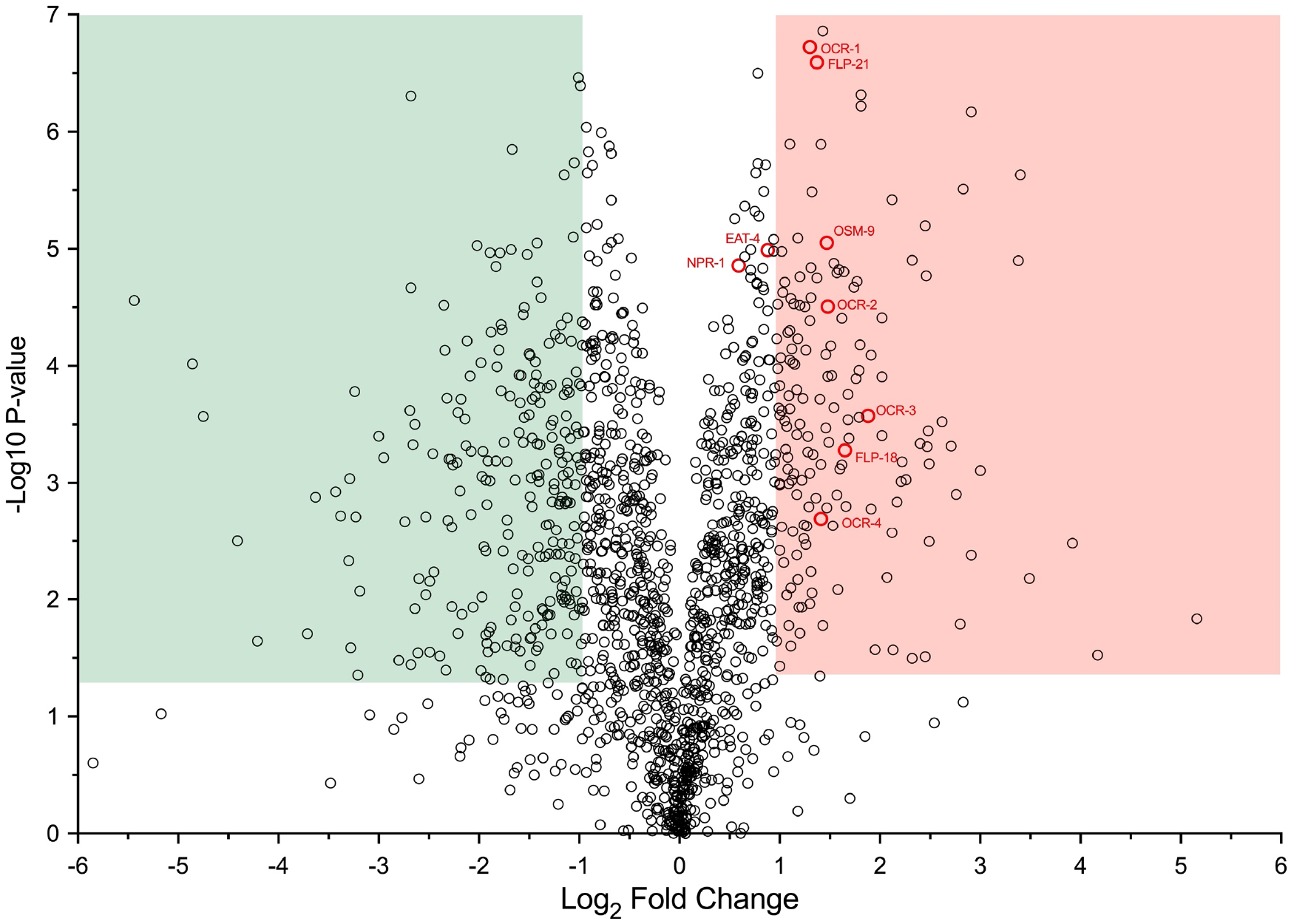
Volcano plot illustrating differential abundances of proteins with the x-axis showing the log2 ratio with respect to the control (N2) and the y-axis representing the −log10 of the p-value. Boxes illustrate a p-value of 0.05, and the positions of the 1.0 and −1.0 log2 ratios that correspond to twofold increases/decreases are colored green and red, respectively. A total of 157 upregulated and 305 downregulated proteins were observed. Interestingly, vanilloid receptor proteins (OCR1-4 and OSM-9) and specific proneuropeptides (FPL-21 and FPL-18) were significantly upregulated following 60 min of exposure to RTX.

Gene Ontology (GO) term enrichment analysis was performed using Metascape [31], and label-free proteomic results included all differentially expressed proteins (DEPs). The top enriched terms are shown in Figure 5A. GO annotation indicated that the differentially expressed proteins were mainly enriched in cellular energy processes, cellular component organization, biosynthesis of amino acids and peptide synthesis and metabolic processes. Moreover, cellular homeostasis is a process associated with ion homeostasis (Ca^2+^, K^+^, Cl^−^ and others). This process is essential to maintain a steady state at the cellular level. RTX is a pungent TRPV1 ligand and activates inward currents due to calcium influx, resulting in cell depolarization. Calcium ion accumulation can provoke apoptosis, and this is the mechanism at the heart of chemical denervation using the agonistic effects of RTX on TRPV1 [4,7]. Interestingly, apoptosis or cell death terms were not among the top enriched GO terms. This is certainly compatible with the behavior results showing that the nematodes completely recovered noxious heat sensitivity at 6 h after RTX exposure, as shown in Figure 2. Consequently, RTX did not provoke nociceptive neuronal death. Kyoto Encyclopedia of Genes and Genomes (KEGG) and REACTOME pathway analyses were performed using ClueGO including all DEPs [32]. The top pathway enrichments are revealed in Figure 5B. As expected, the results were compatible with the GO term enrichment results, with an important emphasis on cellular energy processes and metabolism. ClueGO identified all statistically enriched pathways, and cumulative hypergeometric p-values and enrichment factors were calculated and used for filtering. Selected significant pathways were clustered hierarchically into an interactome created based on Kappa-statistical similarities among their genes (i.e., encoding protein) memberships. Subsequently, a kappa score of 0.8 was applied as the threshold to produce interactome and pathway clusters. The network analysis and interaction between the enriched KEGG and REACTOME pathways are shown in Figure 6. Interestingly, we observed an interaction between the cellular response to chemical stress, beta-catenin-independent Wnt signaling and ion homeostasis. This clearly suggests that these processes are associated with the pungent effects of RTX on the *C. elegans* vanilloid receptor, most likely OCR-3, as shown in Figure 3. The interaction with metabolic processes seems to suggest that they will trigger cellular energy processes involved in catabolism and anabolism reactions. Severe metabolic stress can lead to a state of cell survival, and notably, energy metabolism processes can rescue the cell from apoptosis. However, we were particularly interested in the beta-catenin-independent Wnt signaling pathway. Wnt signaling activates several different intracellular pathways essential to cell proliferation, differentiation, and polarity [44]. Moreover, in *C. elegans*, the Wnt signaling pathway regulates the subcellular positioning of presynaptic terminals and is therefore important for the assembly of synapses, a key component in the formation of neural circuits [45]. Dysfunction in key Wnt pathway components can lead to disorders including depression and anxiety [42–44]. This is consistent with our observation that RTX denervation of cardiac sensory afferents led to a significant reduction in depressive behavior in a mouse model of chronic heart failure [7]. During this study, we also demonstrated the importance of the Wnt signaling pathway. The noncanonical beta-catenin-independent Wnt pathway is linked with Ca^2+^ release from intracellular stores, and an increase in intracellular Ca^2+^ concentration can activate the mitogen-activated protein kinase (MAPK) pathway. Subsequently, this pathway will negatively regulate the canonical Wnt pathway by inhibiting gene transcription mediated by the beta-catenin/TCF complex [42]. Interestingly, canonical Wnt pathway antagonists can effectively mediate depressive behavior [43,44].

**Figure 5.**
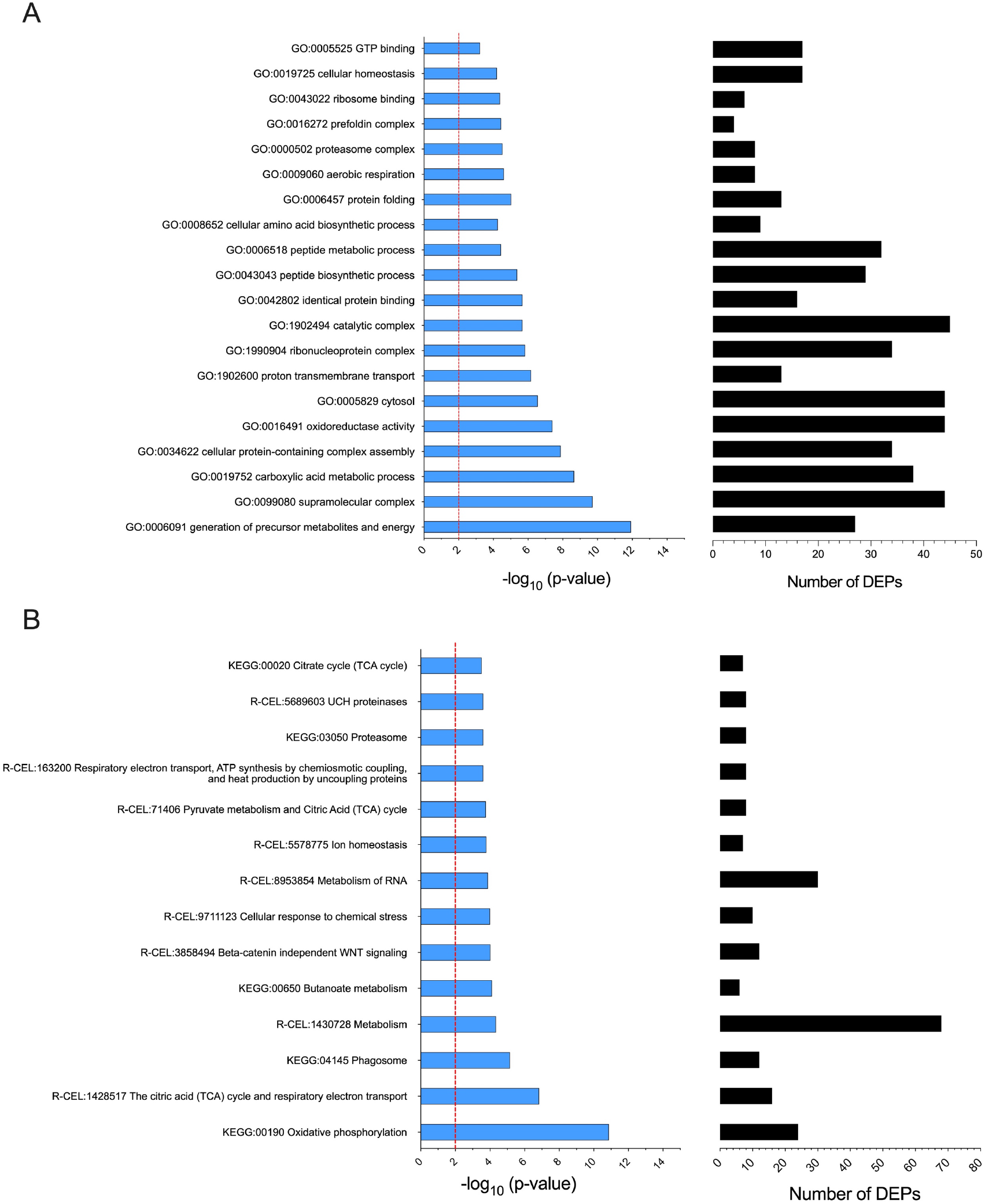
Gene ontology (GO), Kyoto Encyclopedia of Genes and Genomes (KEGG) and REACTOME enriched terms. Analysis was performed with all DEPs (upregulated and downregulated). A) Detailed information relating to changes in the biological processes (BPs), cellular components (CCs), and molecular functions (MFs) of the DEPs following RTX exposure through GO enrichment analyses. B) KEGG and REACTOME pathway enrichment analysis.

**Figure 6.**
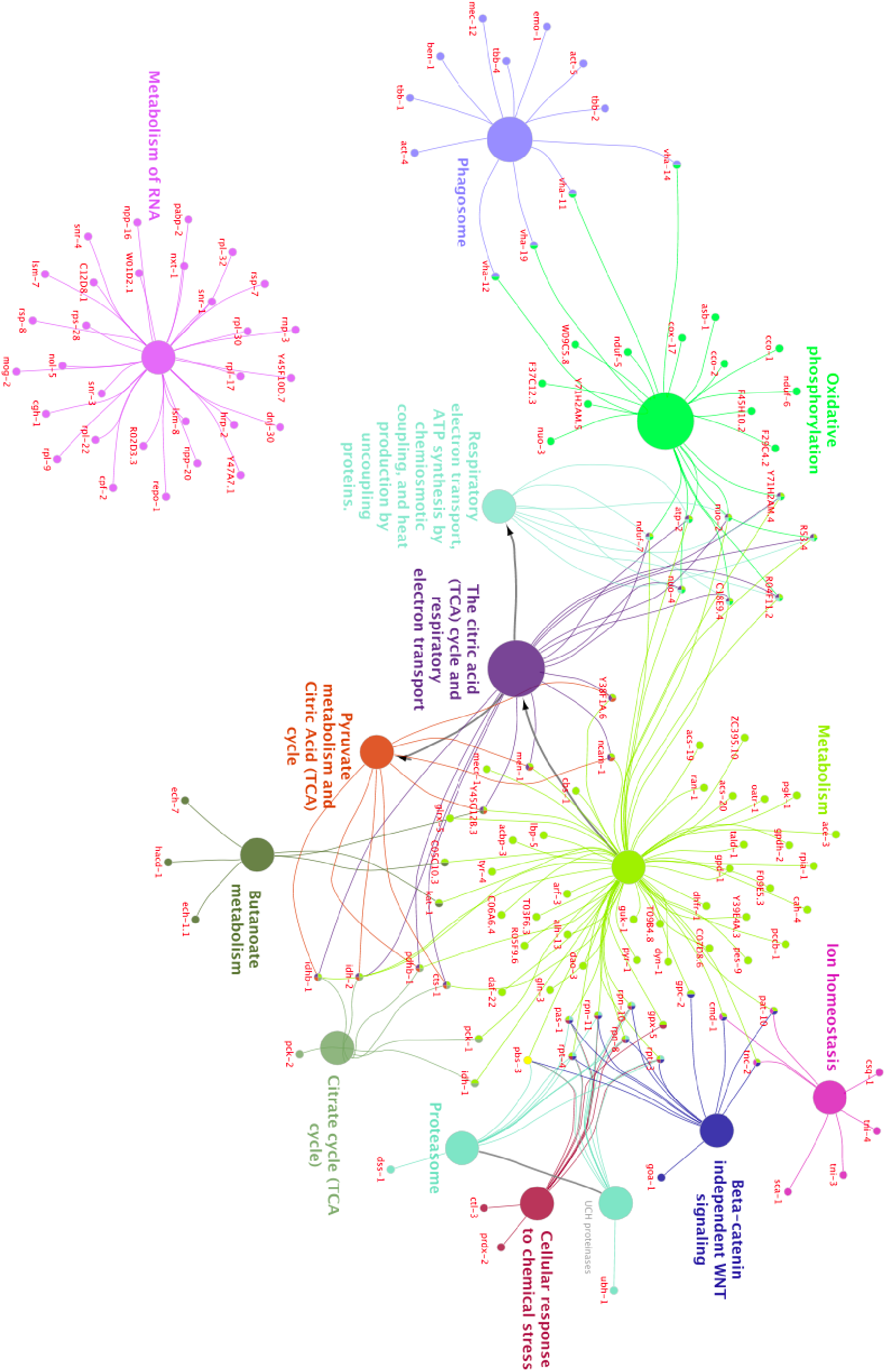
Network analysis and interactions between the enriched KEGG and REACTOME pathways. The network is colored by cluster ID. Please note that nodes sharing the same cluster ID are commonly close to each other. We applied a 0.8 kappa score as the threshold.

## Conclusion

This study has shown for the first time the antinociceptive effects of RTX in *C. elegans* following controlled and prolonged exposure. Moreover, our results suggest that RTX targets the *C. elegans* vanilloid receptor ortholog OCR-3 and not OCR-2, which is the target of capsaicin. Proteomics and bioinformatics analyses revealed that the effects observed with RTX might be related to the noncanonical beta-catenin-independent Wnt pathway leading to negative regulation of the canonical Wnt pathway. This discovery is consistent with our previous study, where we showed that RTX denervation of cardiac sensory afferents yielded a significant reduction in depressive-like behavior in a mouse model of chronic heart failure [7].

## Acknowledgments

This project was funded by the National Sciences and Engineering Research Council of Canada (F. Beaudry discovery grant nos. RGPIN-2015-05071 and RGPIN-2020-05228). Laboratory equipment was funded by the Canadian Foundation for Innovation (CFI) and the *Fonds de Recherche du Québec (FRQ)*, the Government of Quebec (F. Beaudry CFI John R. Evans Leaders grant no. 36706). Ph.D. scholarships were awarded to J. Ben Salem from the Fonds de Recherche du Québec - Santé and from the Université de Montréal.

## Conflicts of interest

The authors declare they have no conflicts of interest.

## Notes

### Competing Interest Statement

The authors have declared no competing interest.

