## Supplemental Data for "Resiniferatoxin hampers the nocifensive response of *Caenorhabditis elegans* to noxious heat, and pathway analysis revealed that the Wnt signaling pathway is involved"

<u>**Supplementary Data**</u>

**Figure S1.** A schematic of the four quadrants assay adapted from Margie et al. (2013). For head avoidance assay, plates were divided into quadrants two test (A and D) and two controls (B and C). Sodium azide was added to all four quadrants to paralyze nematodes. *C. elegans* were added at the center of the plate (typically, n = 100 to 1,000) and after 30 minutes, animals were counted on each quadrant. Only animals outside the inner circle were scored. The calculation of thermal avoidance index was performed has described.

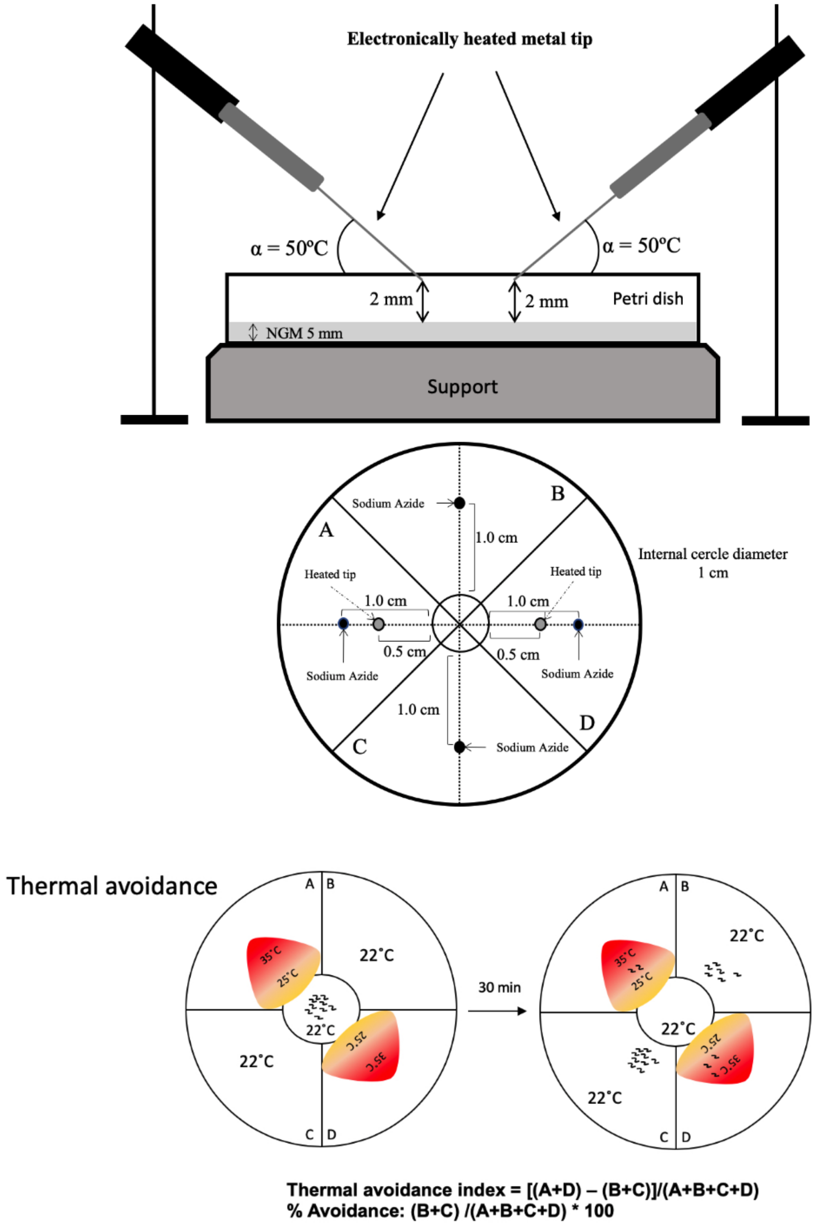

**Table S1**: Data of tested strains and experimental groups described in Figure 1.

| Experimental group | Number of petri dishes | Total number of nematodes counted | Mean % | SD |
|---|---|---|---|---|
| N2(WT) | 12 | 1,241 | 50.87 | 4.02 |
| + RTX 0.1μM | 6 | 994 | 51.31 | 2.92 |
| + RTX 1μM | 8 | 1,795 | 48.99 | 2.87 |
| + RTX 5μM | 8 | 1,438 | 48.54 | 4.82 |

**Table S2**: Data of tested strains and experimental groups described in Figure 2.

| Experimental group | Number of petri dishes | Total number of nematodes counted | Mean % | SD |
|---|---|---|---|---|
| N2(WT) | 12 | 4,566 | 77.29 | 5.03 |
| + Cap 25μM | 12 | 2,982 | 52.75 | 8.32 |
| + RTX 0.1μM | 12 | 6,326 | 61.75 | 7.87 |
| + RTX 1μM | 12 | 3,772 | 54.50 | 6.30 |
| + RTX 5μM | 12 | 2,980 | 55.28 | 4.67 |
| + RTX 0.1μM- 6h latency | 12 | 9,602 | 72.74 | 5.56 |
| + RTX 1μM- 6h latency | 12 | 5,879 | 76.02 | 7.07 |
| + RTX 5μM- 6h latency | 12 | 4,112 | 73.01 | 7.14 |

**Table S3**: Data of tested strains and experimental groups described in Figure 3

| Experimental group | Number of petri dishes | Total number of nematodes counted | Mean % avoidance | SD |
|---|---|---|---|---|
| N2(WT) | 12 | 4,566 | 77.29 | 5.03 |
| N2(WT) + RTX (1μM) | 12 | 3,772 | 54.50 | 6.30 |
| *osm-9(yz6)* | 12 | 8,211 | 62.38 | 4.78 |
| *osm-9(yz6)* + RTX (1μM) | 12 | 9,256 | 46.85 | 3.72 |
| *ocr-2(yz5)* | 12 | 8,723 | 65.21 | 5.69 |
| *ocr-2(yz5)* + RTX (1μM) | 12 | 11,291 | 50.80 | 1.78 |
| *ocr-1(ak46)* | 12 | 4,587 | 68.68 | 9.24 |
| *ocr-1(ak46)* + RTX (1μM) | 12 | 3,657 | 58.25 | 7.57 |
| *ocr-3(ok1559)* | 12 | 3,984 | 69.85 | 5.54 |
| *ocr-3(1559)* + RTX (1μM) | 12 | 4,392 | 65.36 | 3.82 |
| *ocr-4(vs137)* | 12 | 5,862 | 71.00 | 7.62 |
| *ocr-4(vs137)* + RTX (1μM) | 12 | 5,219 | 61.76 | 6.04 |

**Table S5.** Pathway enrichment analysis: List of significantly enriched pathways

| Pathway | p-value* | Coverage (%) | Associated Proteins |
|---|---|---|---|
| Citrate cycle (TCA cycle) | 3.04E-04 | 21.21 | cts-1, idh-1, idh-2, idhb-1, pck-1, pck-2, pdhb-1 |
| Oxidative phosphorylation | 1.36E-11 | 21.24 | C18E9.4, F29C4.2, F37C12.3, F45H10.2, R04F11.2, R53.4, W09C5.8, Y71H2AM.4, Y71H2AM.5, asb-1, atp-2, cco-1, cco-2, cox-17, nduf-5, nduf-6, nduf-7, nuo-2, nuo-3, nuo-4, vha-11, vha-12, vha-14, vha-19 |
| Respiratory electron transportATP synthesis by chemiosmotic coupling, and heat production by uncoupling proteins. | 2.48E-04 | 19.05 | C18E9.4, R04F11.2, R53.4, Y71H2AM.4, atp-2, nduf-7, nuo-2, nuo-4 |
| Beta-catenin independent WNT signaling | 9.70E-05 | 14.81 | cmd-1, goa-1, gpc-2, pas-1, pat-10, pbs-3, rpn-10, rpn-11, rpn-8, rpt-3, rpt-4, tnc-2 |
| Ion homeostasis | 1.61E-04 | 23.33 | cmd-1, csq-1, pat-10, sca-1, tnc-2, tni-3, tni-4 |
| Pyruvate metabolism and Citric Acid (TCA) cycle | 1.74E-04 | 20.00 | Y38F1A.6, Y45G12B.3, cts-1, idh-2, idhb-1, men-1, ncam-1, pdhb-1 |
| Metabolism of RNA | 1.31E-04 | 8.47 | C12D8.1, R02D3.3, W01D2.1, Y45F10D.7, Y47A7.1, cgh-1, cpf-2, dnj-30, hrp-2, lsm-7, lsm-8, mog-2, nol-5, npp-16, npp-20, nxt-1, pabp-2, repo-1, rnp-3, rpl-17, rpl-22, rpl-30, rpl-32, rpl-9, rps-28, rsp-7, rsp-8, snr-1, snr-3, snr-4 |
| Cellular response to chemical stress | 1.02E-04 | 17.24 | ctl-3, gpx-5, pas-1, pbs-3, prdx-2, rpn-10, rpn-11, rpn-8, rpt-3, rpt-4 |
| Butanoate metabolism | 7.68E-05 | 31.58 | C05C10.3, Y45G12B.3, ech-1.1, ech-7, hacd-1, kat-1 |
| Phagosome | 7.05E-06 | 19.05 | act-4, act-5, ben-1, emo-1, mec-12, tbb-1, tbb-2, tbb-4, vha-11, vha-12, vha-14, vha-19 |
| The citric acid (TCA) cycle and respiratory electron transport | 1.42E-07 | 19.51 | C18E9.4, R04F11.2, R53.4, Y38F1A.6, Y45G12B.3, Y71H2AM.4, atp-2, cts-1, idh-2, idhb-1, men-1, ncam-1, nduf-7, nuo-2, nuo-4, pdhb-1 |
| Metabolism | 4.54E-05 | 6.49 | C05C10.3, C06A6.4, C07D8.6, C18E9.4, F09E5.3, R04F11.2, R05F9.6, R53.4, T03F6.3, T09B4.8, Y38F1A.6, Y39E4A.3, Y45G12B.3, Y71H2AM.4, ZC395.10, acbp-3, ace-3, acs-19, acs-20, alh-13, arf-3, |

| | | | |
|---|---|---|---|
| | | | atp-2, cah-4, cbs-1, cmd-1, cts-1, daf-22, dao-3, dhfr-1, dyn-1, gln-3, glrx-5, gpc-2, gpd-1, gpdh-2, gpx-5, guk-1, idh-1, idh-2, idhb-1, kat-1, lbp-5, mecr-1, men-1, ncam-1, nduf-7, nuo-2, nuo-4, oatr-1, pas-1, pat-10, pbs-3, pccb-1, pck-1, pdhb-1, pes-9, pgk-1, pyr-1, ran-1, rpia-1, rpn-10, rpn-11, rpn-8, rpt-3, rpt-4, tald-1, tnc-2, tyr-4 |
| Proteasome | 2.48E-04 | 19.05 | dss-1, pas-1, pbs-3, rpn-10, rpn-11, rpn-8, rpt-3, rpt-4 |
| UCH proteinases | 2.48E-04 | 19.05 | pas-1, pbs-3, rpn-10, rpn-11, rpn-8, rpt-3, rpt-4, ubh-1 |

* p-value was calculated using a right-sided hypergeometric distribution with multiple testing correction (Benjamini & Hochberg (BH) method).

Table S6 All differentially expressed proteins (DEPs) with p-value ≤ 0.01 and used for GO and pathway analyses.

| Ensembl Gene ID **Gene encoding protein** | $log_2$ ratio |
|---|---|
| WBGene00000037 | 5.16 |
| WBGene00013602 | 4.17 |
| WBGene00001248 | 3.92 |
| WBGene00001411 | 3.49 |
| WBGene00019641 | 3.45 |
| WBGene00022745 | 3.4 |
| WBGene00006733 | 3.38 |
| WBGene00001537 | 3 |
| WBGene00044737 | 2.91 |
| WBGene00014016 | 2.91 |
| WBGene00001732 | 2.83 |
| WBGene00009041 | 2.83 |
| WBGene00012165 | 2.8 |
| WBGene00008424 | 2.76 |
| WBGene00021322 | 2.71 |
| WBGene00006776 | 2.62 |
| WBGene00015509 | 2.54 |
| WBGene00009218 | 2.49 |
| WBGene00000474 | 2.49 |
| WBGene00019601 | 2.48 |
| WBGene00007624 | 2.47 |
| WBGene00043057 | 2.46 |
| WBGene00020245 | 2.45 |
| WBGene00019558 | 2.45 |
| WBGene00018385 | 2.4 |
| WBGene00008683 | 2.37 |
| WBGene00000236 | 2.32 |
| WBGene00009854 | 2.32 |
| WBGene00021883 | 2.26 |
| WBGene00022127 | 2.22 |
| WBGene00001214 | 2.21 |
| WBGene00002249 | 2.2 |
| WBGene00020651 | 2.17 |
| WBGene00013869 | 2.13 |
| WBGene00016977 | 2.12 |

| | |
|---|---|
| WBGene00012004 | 2.12 |
| WBGene00004259 | 2.07 |
| WBGene00022276 | 2.02 |
| WBGene00009542 | 2.02 |
| WBGene00007633 | 2.02 |
| WBGene00020832 | 1.97 |
| WBGene00015510 | 1.95 |
| WBGene00021922 | 1.91 |
| WBGene00007010 | 1.91 |
| WBGene00006937 | 1.85 |
| WBGene00018975 | 1.81 |
| WBGene00003802 | 1.81 |
| WBGene00001744 | 1.8 |
| WBGene00007784 | 1.79 |
| WBGene00006047 | 1.79 |
| WBGene00012701 | 1.77 |
| WBGene00004704 | 1.76 |
| WBGene00017484 | 1.74 |
| WBGene00011096 | 1.7 |
| WBGene00001548 | 1.69 |
| WBGene00009778 | 1.68 |
| WBGene00045483 | 1.68 |
| WBGene00023498 | 1.66 |
| WBGene00000371 | 1.64 |
| WBGene00001329 | 1.62 |
| WBGene00019450 | 1.62 |
| WBGene00014178 | 1.6 |
| WBGene00016419 | 1.59 |
| WBGene00011571 | 1.58 |
| WBGene00011977 | 1.57 |
| WBGene00011487 | 1.57 |
| WBGene00011936 | 1.54 |
| WBGene00002060 | 1.54 |
| WBGene00003523 | 1.53 |
| WBGene00001061 | 1.52 |
| WBGene00006997 | 1.51 |
| WBGene00004771 | 1.49 |
| WBGene00013308 | 1.48 |
| WBGene00012179 | 1.47 |

| | |
|---|---|
| WBGene00007458 | 1.46 |
| WBGene00006725 | 1.46 |
| WBGene00016900 | 1.43 |
| WBGene00019572 | 1.43 |
| WBGene00022492 | 1.41 |
| WBGene00013866 | 1.41 |
| WBGene00009031 | 1.4 |
| WBGene00008603 | 1.4 |
| WBGene00001048 | 1.37 |
| WBGene00019821 | 1.36 |
| WBGene00000376 | 1.34 |
| WBGene00012326 | 1.33 |
| WBGene00022615 | 1.32 |
| WBGene00011308 | 1.32 |
| WBGene00018240 | 1.31 |
| WBGene00000931 | 1.31 |
| WBGene00020894 | 1.31 |
| WBGene00001538 | 1.3 |
| WBGene00001113 | 1.3 |
| WBGene00004444 | 1.3 |
| WBGene00020732 | 1.29 |
| WBGene00000402 | 1.28 |
| WBGene00001682 | 1.28 |
| WBGene00000370 | 1.27 |
| WBGene00011880 | 1.26 |
| WBGene00020242 | 1.26 |
| WBGene00004386 | 1.25 |
| WBGene00008210 | 1.24 |
| WBGene00010664 | 1.24 |
| WBGene00004499 | 1.24 |
| WBGene00011156 | 1.23 |
| WBGene00001150 | 1.22 |
| WBGene00007154 | 1.22 |
| WBGene00017374 | 1.2 |
| WBGene00020097 | 1.2 |
| WBGene00017482 | 1.2 |
| WBGene00004466 | 1.2 |
| WBGene00013127 | 1.2 |
| WBGene00011730 | 1.19 |

| | |
|---|---|
| WBGene00009189 | 1.18 |
| WBGene00013077 | 1.18 |
| WBGene00010879 | 1.18 |
| WBGene00013393 | 1.18 |
| WBGene00017609 | 1.17 |
| WBGene00004217 | 1.17 |
| WBGene00008975 | 1.17 |
| WBGene00019042 | 1.17 |
| WBGene00002001 | 1.15 |
| WBGene00006994 | 1.14 |
| WBGene00019719 | 1.14 |
| WBGene00007627 | 1.13 |
| WBGene00014262 | 1.13 |
| WBGene00018237 | 1.12 |
| WBGene00018346 | 1.12 |
| WBGene00018742 | 1.11 |
| WBGene00000465 | 1.11 |
| WBGene00004889 | 1.11 |
| WBGene00020379 | 1.11 |
| WBGene00004138 | 1.1 |
| WBGene00012557 | 1.1 |
| WBGene00020915 | 1.1 |
| WBGene00219313 | 1.1 |
| WBGene00014252 | 1.09 |
| WBGene00010357 | 1.09 |
| WBGene00021751 | 1.09 |
| WBGene00021024 | 1.09 |
| WBGene00013063 | 1.08 |
| WBGene00013064 | 1.08 |
| WBGene00004701 | 1.08 |
| WBGene00011536 | 1.08 |
| WBGene00013867 | 1.08 |
| WBGene00010631 | 1.07 |
| WBGene00018159 | 1.07 |
| WBGene00022170 | 1.06 |
| WBGene00006038 | 1.05 |
| WBGene00006040 | 1.05 |
| WBGene00004497 | 1.04 |
| WBGene00004446 | 1.03 |

| | |
|---|---|
| WBGene00016658 | 1.02 |
| WBGene00000928 | 1.02 |
| WBGene00019261 | 1.02 |
| WBGene00007534 | 1.01 |
| WBGene00001604 | 1 |
| WBGene00013683 | 1 |
| WBGene00003390 | -1 |
| WBGene00003806 | -1 |
| WBGene00018460 | -1.01 |
| WBGene00018461 | -1.01 |
| WBGene00018151 | -1.02 |
| WBGene00010768 | -1.02 |
| WBGene00003082 | -1.02 |
| WBGene00006826 | -1.03 |
| WBGene00000534 | -1.03 |
| WBGene00003916 | -1.03 |
| WBGene00020185 | -1.03 |
| WBGene00004434 | -1.03 |
| WBGene00019890 | -1.03 |
| WBGene00011292 | -1.04 |
| WBGene00007071 | -1.04 |
| WBGene00016050 | -1.04 |
| WBGene00000206 | -1.05 |
| WBGene00001000 | -1.05 |
| WBGene00006730 | -1.05 |
| WBGene00003762 | -1.05 |
| WBGene00008452 | -1.06 |
| WBGene00000067 | -1.08 |
| WBGene00017719 | -1.08 |
| WBGene00000710 | -1.08 |
| WBGene00002183 | -1.08 |
| WBGene00007110 | -1.09 |
| WBGene00017926 | -1.09 |
| WBGene00016316 | -1.09 |
| WBGene00004503 | -1.1 |
| WBGene00006586 | -1.1 |
| WBGene00011938 | -1.1 |
| WBGene00007420 | -1.1 |
| WBGene00012999 | -1.1 |

| | |
|---|---|
| WBGene00019962 | -1.1 |
| WBGene00011637 | -1.11 |
| WBGene00018610 | -1.11 |
| WBGene00009394 | -1.12 |
| WBGene00008591 | -1.12 |
| WBGene00021952 | -1.12 |
| WBGene00018701 | -1.12 |
| WBGene00018145 | -1.12 |
| WBGene00015514 | -1.12 |
| WBGene00009818 | -1.12 |
| WBGene00006921 | -1.13 |
| WBGene00022182 | -1.13 |
| WBGene00005000 | -1.13 |
| WBGene00004736 | -1.13 |
| WBGene00012713 | -1.13 |
| WBGene00004491 | -1.14 |
| WBGene00004302 | -1.14 |
| WBGene00020369 | -1.14 |
| WBGene00016333 | -1.14 |
| WBGene00017657 | -1.14 |
| WBGene00009092 | -1.14 |
| WBGene00021839 | -1.15 |
| WBGene00011018 | -1.15 |
| WBGene00004467 | -1.15 |
| WBGene00003234 | -1.16 |
| WBGene00021613 | -1.16 |
| WBGene00018271 | -1.17 |
| WBGene00004420 | -1.17 |
| WBGene00010130 | -1.17 |
| WBGene00009514 | -1.17 |
| WBGene00020781 | -1.17 |
| WBGene00003989 | -1.18 |
| WBGene00000292 | -1.18 |
| WBGene00010094 | -1.18 |
| WBGene00022173 | -1.19 |
| WBGene00000552 | -1.19 |
| WBGene00002254 | -1.19 |
| WBGene00000833 | -1.2 |
| WBGene00004410 | -1.2 |

| | |
|---|---|
| WBGene00020931 | -1.2 |
| WBGene00012887 | -1.21 |
| WBGene00011273 | -1.22 |
| WBGene00000973 | -1.23 |
| WBGene00007942 | -1.23 |
| WBGene00001088 | -1.24 |
| WBGene00016524 | -1.24 |
| WBGene00009739 | -1.24 |
| WBGene00004705 | -1.25 |
| WBGene00003994 | -1.25 |
| WBGene00013220 | -1.25 |
| WBGene00008422 | -1.25 |
| WBGene00010303 | -1.26 |
| WBGene00005007 | -1.26 |
| WBGene00202498 | -1.27 |
| WBGene00000215 | -1.27 |
| WBGene00022497 | -1.27 |
| WBGene00021826 | -1.28 |
| WBGene00002272 | -1.28 |
| WBGene00018811 | -1.28 |
| WBGene00008505 | -1.29 |
| WBGene00000786 | -1.29 |
| WBGene00002008 | -1.3 |
| WBGene00009362 | -1.3 |
| WBGene00000479 | -1.3 |
| WBGene00008335 | -1.31 |
| WBGene00015192 | -1.31 |
| WBGene00020391 | -1.32 |
| WBGene00000229 | -1.32 |
| WBGene00006920 | -1.33 |
| WBGene00011232 | -1.33 |
| WBGene00010673 | -1.34 |
| WBGene00017283 | -1.36 |
| WBGene00000293 | -1.36 |
| WBGene00007684 | -1.37 |
| WBGene00022230 | -1.37 |
| WBGene00000991 | -1.38 |
| WBGene00001748 | -1.39 |
| WBGene00006486 | -1.39 |

| | |
|---|---|
| WBGene00006434 | -1.39 |
| WBGene00021636 | -1.39 |
| WBGene00015565 | -1.4 |
| WBGene00021043 | -1.4 |
| WBGene00001694 | -1.41 |
| WBGene00012185 | -1.41 |
| WBGene00019205 | -1.42 |
| WBGene00004464 | -1.42 |
| WBGene00009367 | -1.42 |
| WBGene00012376 | -1.42 |
| WBGene00004917 | -1.43 |
| WBGene00003836 | -1.43 |
| WBGene00010605 | -1.44 |
| WBGene00008183 | -1.44 |
| WBGene00012375 | -1.44 |
| WBGene00011155 | -1.45 |
| WBGene00011399 | -1.45 |
| WBGene00000929 | -1.47 |
| WBGene00003081 | -1.47 |
| WBGene00017075 | -1.47 |
| WBGene00019178 | -1.47 |
| WBGene00015752 | -1.48 |
| WBGene00010117 | -1.48 |
| WBGene00000780 | -1.48 |
| WBGene00010041 | -1.49 |
| WBGene00001303 | -1.49 |
| WBGene00001030 | -1.49 |
| WBGene00012344 | -1.49 |
| WBGene00002257 | -1.49 |
| WBGene00018024 | -1.5 |
| WBGene00000248 | -1.5 |
| WBGene00021647 | -1.5 |
| WBGene00021934 | -1.51 |
| WBGene00003143 | -1.51 |
| WBGene00002273 | -1.51 |
| WBGene00002978 | -1.52 |
| WBGene00004504 | -1.52 |
| WBGene00010317 | -1.54 |
| WBGene00004429 | -1.55 |

| | |
|---|---|
| WBGene00015185 | -1.55 |
| WBGene00016435 | -1.56 |
| WBGene00000469 | -1.56 |
| WBGene00020833 | -1.56 |
| WBGene00010266 | -1.56 |
| WBGene00001784 | -1.57 |
| WBGene00000214 | -1.58 |
| WBGene00006824 | -1.58 |
| WBGene00004439 | -1.59 |
| WBGene00012608 | -1.59 |
| WBGene00017184 | -1.6 |
| WBGene00001500 | -1.6 |
| WBGene00000183 | -1.61 |
| WBGene00001498 | -1.62 |
| WBGene00009051 | -1.62 |
| WBGene00009454 | -1.63 |
| WBGene00009625 | -1.64 |
| WBGene00008378 | -1.64 |
| WBGene00009583 | -1.64 |
| WBGene00007993 | -1.65 |
| WBGene00001648 | -1.66 |
| WBGene00001501 | -1.67 |
| WBGene00008053 | -1.68 |
| WBGene00002000 | -1.68 |
| WBGene00007974 | -1.69 |
| WBGene00019783 | -1.69 |
| WBGene00000182 | -1.7 |
| WBGene00003922 | -1.71 |
| WBGene00010556 | -1.71 |
| WBGene00010778 | -1.72 |
| WBGene00000170 | -1.72 |
| WBGene00006889 | -1.72 |
| WBGene00012354 | -1.75 |
| WBGene00009987 | -1.75 |
| WBGene00002269 | -1.76 |
| WBGene00006721 | -1.76 |
| WBGene00012983 | -1.77 |
| WBGene00022499 | -1.77 |
| WBGene00003903 | -1.78 |

| | |
|---|---|
| WBGene00021564 | -1.78 |
| WBGene00003982 | -1.78 |
| WBGene00002065 | -1.79 |
| WBGene00004111 | -1.79 |
| WBGene00007969 | -1.8 |
| WBGene00022169 | -1.81 |
| WBGene00018218 | -1.81 |
| WBGene00019727 | -1.82 |
| WBGene00012964 | -1.83 |
| WBGene00006706 | -1.83 |
| WBGene00001683 | -1.84 |
| WBGene00011330 | -1.86 |
| WBGene00021204 | -1.88 |
| WBGene00004201 | -1.88 |
| WBGene00003904 | -1.89 |
| WBGene00019978 | -1.89 |
| WBGene00002244 | -1.89 |
| WBGene00006538 | -1.9 |
| WBGene00020339 | -1.91 |
| WBGene00007646 | -1.91 |
| WBGene00006460 | -1.92 |
| WBGene00015814 | -1.92 |
| WBGene00007653 | -1.93 |
| WBGene00007258 | -1.93 |
| WBGene00012553 | -1.93 |
| WBGene00010845 | -1.93 |
| WBGene00020192 | -1.94 |
| WBGene00020417 | -1.95 |
| WBGene00003175 | -1.96 |
| WBGene00020382 | -1.97 |
| WBGene00019401 | -1.97 |
| WBGene00014030 | -1.97 |
| WBGene00003949 | -1.98 |
| WBGene00000282 | -1.98 |
| WBGene00006565 | -2 |
| WBGene00008218 | -2.02 |
| WBGene00008607 | -2.06 |
| WBGene00013284 | -2.08 |
| WBGene00008334 | -2.08 |

| | |
|---|---|
| WBGene00003370 | -2.09 |
| WBGene00013029 | -2.1 |
| WBGene00007927 | -2.12 |
| WBGene00001130 | -2.13 |
| WBGene00009188 | -2.14 |
| WBGene00000066 | -2.17 |
| WBGene00019220 | -2.18 |
| WBGene00003230 | -2.18 |
| WBGene00013227 | -2.19 |
| WBGene00020588 | -2.19 |
| WBGene00000822 | -2.21 |
| WBGene00023450 | -2.21 |
| WBGene00012932 | -2.22 |
| WBGene00004914 | -2.26 |
| WBGene00003934 | -2.27 |
| WBGene00019179 | -2.27 |
| WBGene00017023 | -2.28 |
| WBGene00006537 | -2.28 |
| WBGene00020335 | -2.3 |
| WBGene00007516 | -2.31 |
| WBGene00020190 | -2.32 |
| WBGene00001005 | -2.32 |
| WBGene00006583 | -2.32 |
| WBGene00003968 | -2.32 |
| WBGene00001776 | -2.33 |
| WBGene00006536 | -2.34 |
| WBGene00014108 | -2.35 |
| WBGene00019682 | -2.39 |
| WBGene00000774 | -2.41 |
| WBGene00001037 | -2.45 |
| WBGene00010759 | -2.46 |
| WBGene00006585 | -2.46 |
| WBGene00009004 | -2.47 |
| WBGene00000435 | -2.49 |
| WBGene00011736 | -2.49 |
| WBGene00003522 | -2.51 |
| WBGene00016953 | -2.53 |
| WBGene00016422 | -2.53 |
| WBGene00019324 | -2.6 |

| | |
|---|---|
| WBGene00001233 | -2.6 |
| WBGene00013924 | -2.61 |
| WBGene00021888 | -2.64 |
| WBGene00022435 | -2.64 |
| WBGene00004387 | -2.66 |
| WBGene00001156 | -2.68 |
| WBGene00022267 | -2.68 |
| WBGene00004916 | -2.68 |
| WBGene00013639 | -2.69 |
| WBGene00021657 | -2.74 |
| WBGene00022599 | -2.77 |
| WBGene00020717 | -2.8 |
| WBGene00015413 | -2.85 |
| WBGene00004145 | -2.9 |
| WBGene00002064 | -2.95 |
| WBGene00011275 | -3 |
| WBGene00015101 | -3.09 |
| WBGene00009995 | -3.19 |
| WBGene00021465 | -3.21 |
| WBGene00008339 | -3.23 |
| WBGene00021286 | -3.24 |
| WBGene00011015 | -3.28 |
| WBGene00016594 | -3.29 |
| WBGene00004172 | -3.3 |
| WBGene00010796 | -3.38 |
| WBGene00018393 | -3.38 |
| WBGene00021158 | -3.43 |
| WBGene00020112 | -3.48 |
| WBGene00001854 | -3.63 |
| WBGene00004986 | -3.67 |
| WBGene00007330 | -3.71 |
| WBGene00021930 | -4.11 |
| WBGene00002149 | -4.21 |
| WBGene00006508 | -4.41 |
